# The effect of dietary chitin on Atlantic salmon (*Salmo salar*) chitinase activity, gene expression, and microbial composition

**DOI:** 10.1101/2022.05.05.490722

**Authors:** Matilde Mengkrog Holen, Simen Rød Sandve, Thomas Nelson Harvey, Yang Jin, Inga Leena Angell, Knut Rudi, Matthew Peter Kent

## Abstract

**Background:** Chitin is a common component in the natural diet of many fish, and a range of chitinases with the potential to down chitin have been identified. Yet whether chitin is metabolized in fish is still unclear. Here we used a combination of chitinase activity assay, transcriptomics, and 16S rRNA bacterial analysis to assess the effect of chitin supplementation on Atlantic salmon gene expression and microbial community.

**Results:** Atlantic salmon express multiple genes associated with chitin metabolism, and we show that the expression and activity of Atlantic salmon chitinases are not affected by the addition of dietary chitin. We do, however, demonstrate an association between gut microbial composition, chitinase activity in the gut, and host chitinase expression.

**Conclusion:** The findings presented here support the idea that chitin metabolism genes are linked to the maintenance of a chitin-based barrier in the teleost gut. These results contribute to a greater understanding of chitin metabolism in fish.

## INTRODUCTION

Chitin is a tough and insoluble polymer consisting of β-1,4-linked *N*-acetyl-D-glucosamine (GlcNAc) residues that serves as a protective and structural component in arthropod exoskeletons such as insects and crustaceans (Kramer *et al*. 1995; Kurita 2006). This complex carbohydrate is commonly found in the natural diet of many fish, including Atlantic salmon (*Salmo salar*), and chitinous organisms like Antarctic krill (*Euphausia superba*) and black soldier fly (*Hermetia illucens*) are considered two sustainable feed sources for Atlantic salmon farming (Hansen *et al*. 2010; Belghit *et al*. 2019). Supplementation of chitin in Atlantic salmon feed has been shown to promote the growth of potentially beneficial gut microbes that *in vitro* inhibit the growth of common fish pathogens (Askarian *et al*. 2012). However, some studies have reported a reduced growth rate in salmon fed chitin-rich diets and it has been hypothesized that chitin acts as an energy sink when fish are not able to digest and utilize this polysaccharide properly (Karlsen *et al*. 2017; Renna *et al*. 2017).

Chitinase and chitobiase are the enzymes responsible for the hydrolysis of chitin and the production of GlcNAc monomers. Their activity has been detected in the digestive tract of a variety of fish species (Jeuniaux 1961; Fänge *et al*. 1979; Lindsay *et al*. 1984; Lindsay 1984; Kono *et al*. 1987; Gutowska *et al*. 2004; Krogdahl *et al*. 2005; Abro *et al*. 2014), including salmonids such as rainbow trout (*Oncorhynchus mykiss*) (Lindsay *et al*. 1984), and a range of fish chitinases have been identified and biochemically characterized (Ikeda *et al*. 2009, 2012, 2013, 2017; Zhang *et al*. 2012; Koch *et al*. 2014; Teng *et al*. 2014; Kakizaki *et al*. 2015; Kawashima *et al*. 2016; Pohls *et al*. 2016; Gao *et al*. 2017). Most of these enzymes are detected in the gastrointestinal tract and show acid-resistant activities toward insoluble chitin substrates, indicating that they could have the ability to digest chitin (Ikeda *et al*. 2017). As far as we know, no one has characterized chitinase proteins in Atlantic salmon.

Another family of enzymes involved in chitin metabolism is chitin synthases (CHS), which synthesize chitin from GlcNAc. Since chitin is present in their scales (Tang *et al*. 2015), it is hypothesized that salmon express CHS and that these are active, however very little is known about the role of these enzymes in metabolic processes. In this study we investigated the effect of chitin supplementation on the expression of host chitinases, chitobiase, and CHS as well as on the microbiome of Atlantic salmon.

## MATERIALS AND METHODS

### Feeding trial

The fish used in this experiment were Atlantic salmon post-smolts obtained as fry from AquaGen Breeding Centre, Kyrksæterøra, and reared in fresh water at the Centre for Fish Research, NMBU. The fish (n = 32) were acclimatized to experimental conditions for 41 days before the trial and were fed a standard commercial diet. Groups of six fish were placed in each tank with an average weight of 812 ± 113 g and diets were tested in two replicate tanks. The fish were fed one of three different diets (Table 1) over 29 days through an automatic feeding system. Starting with a standard core feed composition, Diet 1 – “control” was supplemented with 6 % cellulose bought from FôrTek, NMBU, Diet 2 – “shrimp shell” was supplemented with 6 % shrimp (*Pandalus borealis*) shell chitin bought from Primex (ChitoClear Chitin, Iceland) and in Diet 3, a portion of the fish meal was substituted with insect meal from defatted black soldier fly larvae (*Hermetica illucens*) obtained from Protix Biosystems BV (Dongen, Netherlands) to a final concentration of approximately 6% chitin. After 29 days fish were euthanized by a blow to the head in accordance with the national regulations of animal (Dyrevelferdsloven 2015). Tissue samples and the contents from the stomach and pyloric caeca were collected separately, flash-frozen in liquid nitrogen, and stored at -80 °C until analysis of gene expression and chitinase activity. Contents from the distal intestine (DI), the most distal compartment of the gut, were sampled for 16S rRNA sequencing as previously described (Rudi *et al*. 2018).

**Table 1.** Ingredients and proximal composition of the experimental diets. **\*** This is a proximate measure using the following formula to calculate chitin in insect meal: chitin (%) 76 = ash-free ADF (%) - ADIP (%) as previously done by Marono S. *et al* (2015) (Marono *et al*. 2015)

|  | *Experimental diets* |  |  |
| --- | --- | --- | --- |
|  | *Control* | *Shrimp shell* | *Fly larvae* |
| *Ingredients (g kg<sup>-1</sup>)* |  |  |  |
| Cellulose | **4.7** | 0.0 | 0.0 |
| Fish meal | 32.2 | 32.2 | 9.1 |
| Wheat gluten | 9.4 | 9.4 | 9.4 |
| Pregel potato starch | 8.6 | 8.6 | 8.6 |
| Defatted black soldier fly larvae meal | 0.0 | 0.0 | **32.4** |
| Shrimp shell chitin | 0.0 | **4.7** | 0.0 |
| Gelatine | 5.8 | 5.8 | 5.8 |
| Choline chloride | 0.1 | 0.1 | 0.1 |
| Fish oil | 10.8 | 10.8 | 6.1 |
| Vitamin/mineral mixture | 0.5 | 0.5 | 0.5 |
| *Proximate composition (%)* |  |  |  |
| Ash | 9.17 | 8.79 | 6.38 |
| Moisture | 6.91 | 6.58 | 4.71 |
| Lipid | 19.64 | 19.64 | 19.25 |
| Crude protein (N · 6.25) | 46.9 | 46.9 | 47.36 |
| Chitin | **0** | **6.11** | **6.35\*** |
| Cellulose | 6.11 | 0 | 0 |

\* This is a proximate measure using the following formula to calculate chitin in insect meal: chitin (%) = ash-free ADF (%) - ADIP (%) as previously done by Marono S. *et al.* (2015) (Marono *et al.* 2015)

### RNA-sequencing

Total RNA was extracted from the stomach and pyloric caeca tissue samples stored at -80 °C (n = 4 for control and fly larvae, n = 5 for shrimp shell) using the RNeasy Plus Universal Kit (QIAGEN). RNA quality was assessed using a 2100 Bioanalyzer with the RNA 6000 nano kit (Agilent) and the concentration was measured using a Nanodrop 8000 spectrophotometer (Thermo Fisher Scientific). Extracted RNA with RNA integrity number (RIN) ≥7 was used as input for the TruSeq Stranded mRNA HT Sample Prep Kit (Illumina) according to the manufacturer ‘s recommendations. Mean library length was measured on 2100 Bioanalyzer using the DNA 1000 kit (Agilent) and the library concentration was quantified with the Qbit Broad Range kit (Thermo Fisher Scientific). Single-end sequencing (100bp reads) was performed at the Norwegian Sequencing Center (Oslo, Norway) using a HiSeq4000 instrument (Illumina).

### Gene expression

Trimming, mapping, and counting of reads were done using the bcbio-nextgen pipeline (https://github.com/bcbio/bcbio-nextgen). The sequencing reads were aligned to the Atlantic salmon genome (ICSASG_v2) (Lien *et al*. 2016) using the STAR aligner with default settings (Dobin *et al*. 2013) after adapter trimming. FeatureCounts was then used to count reads aligned to genes (Liao *et al*. 2014). Raw gene counts were transformed to transcripts per million (TPM) values to normalize for gene length before comparison of chitinase gene expression levels (Welch ‘s t-test, α = 0.05). Gene expression values were normalized by library size (see TMM normalization in EdgeR user guide (Robinson and Oshlack 2010)) before differential expression analysis. Based on PCA analysis, two samples (X8.CFE.7.F4.PC and X9.CFE.13.F1.PC) showed unusual gene expression patterns and were removed from further analysis of pyloric caeca for fish fed control and fly larvae. All statistical analysis was done in R (v.3.6.0).

### Differential expression analysis

Lowly expressed genes (log_2_(TPM + 1) values < 1) were filtered out prior to differential expression analysis. The analysis was carried out using the standard EdgeR protocol (Robinson *et al*. 2010) where an exact test of expression values between the experimental diet (shrimp shell or fly larvae) and control diet gave a log_2_-fold change, p-value, and false discovery rate (FDR) for each gene. Genes with an FDR < 0.05 and absolute log_2_ fold change (logFC) > 0.5 were defined as differentially expressed.

### Gene enrichment analysis

Gene enrichment analysis of the differential expressed genes was performed using KEGG (Kyoto Encyclopedia of Genes and Genomes) and GO (Gene Ontology). KEGG analysis was carried out using the “kegga” function in the limma-package (Ritchie *et al*. 2015) with the argument “species.KEGG = “sasa”“ and p-values < 0.05. GO analysis was carried out using the package GOstats (Falcon and Gentleman 2007), following the steps outlined here (https://www.bioconductor.org/packages/release/bioc/vignettes/GOstats/inst/doc/GOstatsForUnsupportedOrganisms.pdf), using the argument “ontology = “BP”“ (biological processes) and Bonferroni adjusted p-values (q) < 0.05.

### Chitinase activity test

We measured the chitinase activity of fish analyzed for gene expression that contained stomach and pyloric caeca content (n = 3) as described previously (Ohno *et al*. 2013). Approximately 200 mg of contents were homogenized using TissueLyser II (QIAGEN) in 900 uL of 50 mM sodium acetate (pH 124 = 5.5) containing 1X Halt protease inhibitor cocktail (Thermo Fisher Scientific). The samples were centrifuged at 14.000 g for 20 min at 4°C to pellet particulates, and the supernatant was collected for protein quantification and chitinase activity. Protein concentration was measured using Bradford protein assay (Quick Start™ Bradford Assay, BioRad) with BSA (Bovine Serum Albumin) used as standard, according to manufacturer ‘s instructions. Chitinase activity was measured by monitoring the hydrolysis of 4-Methylumbelliferyl β-D-N,N′,N′′-triacetylchitotrioside (Sigma-Aldrich, M5639), a fluorogenic chitin substrate suitable for measuring endochitinase activity, according to the Fluorimetric Chitinase Assay Kit (Sigma-Aldrich, CS1030) with the following modifications. Briefly, 10 µg of pyloric caeca proteins and 1 µg of stomach proteins were incubated with 200 μM 4-Methylumbelliferyl β-D-N,N′,N′′- triacetylchitotrioside (4-MU-GlcNAc_3_) in McIlvaine ‘s buffer (0.1 M citric acid and 0.2 M Na_2_HPO_4_, pH 6) in a volume of 100 µL at 28 °C. The pH and temperature were determined from pilot experiments on a recombinant Atlantic salmon chitinase (rChia.8, unpublished) where pH 6 and 28 °C were optimal conditions for activity of this enzyme which is expressed in both stomach and pyloric caeca of Atlantic salmon. The reaction was terminated after 30 min by adding 400 mM sodium carbonate and the fluorescence of the released 4-Methylumbelliferone (4-MU) was measured at an excitation wavelength of 360 nm and an emission wavelength of 450 nm using a SpectraMax M2 plate reader (Molecular Devices) no later than 30 minutes after stopping the reaction. The assay was performed in triplicates and a 4-MU standard curve was used to quantify 4-MU resulting from the hydrolytic reaction. The measured fluorescence was corrected for hydrolysis of the substrate without enzyme added. The cchitinase activity was expressed as unit/mg of total protein in the sample where 1 unit is defined as 1 umol of 4-MU formed per minute. Statistical comparisons were done using Student ‘s t-test (α = 0.05 145 and α = 0.1).

### Illumina sequencing of the 16S rRNA gene

DNA was extracted from ground samples of Diet 1-3 fish feed (n=2 per feed) and the contents collected from the distal intestine (DI) from all fish (n=12 per feed) using the Mag midi kit (LGC Genomics, UK) following the manufactures instructions. Preparation of amplicon library using the primer pair 341F/806R (Yu *et al*. 2005) and sequencing was done as previously described (Rudi *et al*. 2018). The resulting reads were processed as previously described (Angell *et al*. 2020) using a sequence depth of 10,000 sequences per sample.

## RESULTS

### Impact of dietary chitin on chitinase activity and expression

We performed an *in vitro* quantification of chitinase activity from crude extracts of total soluble material collected from the stomach and pyloric caeca of fish fed one of three diets differing in chitin content. Chitinase activity relative to the amount of protein was consistently much higher (ranging from 5 to11- fold difference) in the stomach than in pyloric caeca irrespective of diets. Chitinase activity was unaffected by the inclusion of dietary chitin when compared to control (Student ‘s t-test, p > 0.05; Figure 1). However, a trend towards lower activity in fish fed fly larvae compared to the fish fed control and shrimp shell was observed (Student ‘s t-test, p < 0.1).

**Figure 1.**
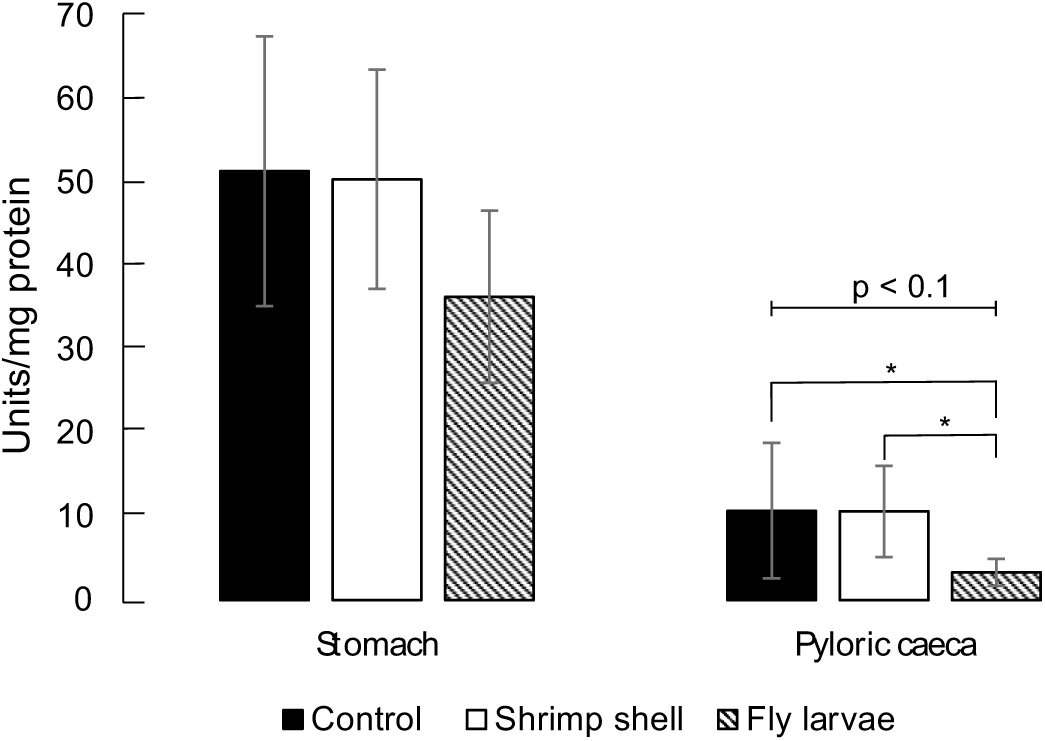
Chitinase activity (units/mg protein) in the stomach- and pyloric caeca contents of fish (n=3) fed control, shrimp shell, and fly larvae using 4-MU-GlcNAc3. Crude soluble protein solutions from stomach (1 μg per reaction) and pyloric caeca (10 μg per reaction) was incubated with 200 μM 4-MU-GlcNAc_3_ for 30 minutes at 28 °C in McIlvaine ‘s buffer, pH 6. 1 unit is defined as 1 µmol of 4-MU formed per minute. The values shown are means of triplicates ± SD.

According to the NCBI Salmo Salar RefSeq annotation (GCF_000233375.1; release 100), Atlantic salmon codes for 10 chitinase-like genes from the glycosyl hydrolase family 18 (hereafter named *chia.1*-*10*; see Supplementary Table 1 for the corresponding gene IDs), a single chitobiase-like gene from the glycosyl hydrolase family 18 (hereafter named *ctbs*), and four genes with putative chitin- synthase domains (hereafter named *chs1a, chs1b, chs2*, and *chs3*). Eight chitinase genes and two CHS genes showed tissue-specific expression (stomach; *chia.3, 4, 5* and *7*, pyloric caeca; *chia.1, 2, 6, 9* and *10, chs1a* and *chs1b*) with *chia.8* and *ctbs* being ubiquitously expressed in both tissues (Figure 2A). Notably, three of the chitinase genes expressed in the stomach (*chia.3, 4*, and *7*) were among the most abundant transcripts present in the tissue, whereas *chia.1, chia.2*, and *chia.6* were among the most abundant transcripts in pyloric caeca. Two CHS genes, *chs2* and *chs3* were not expressed in any of the two tissues. To assess the response of chitinase-, chitobiase- and CHS genes to the inclusion of dietary chitin, we compared their expression levels in the stomach (Figure 2B) and pyloric caeca (Figure 2C). One chitinase, *chia.5*, was lowly expressed (log_2_(TPM) < 2) and is therefore not shown in Figure 2B. The inclusion of dietary chitin had no significant effect on chitinase, chitobiase, and CHS gene expression (Welch ‘s t-test, p > 0.05), although the expression of *chia.1, chia.2*, and *chia.6* in pyloric caeca appeared to be slightly lower in fish fed fly larvae (p < 0.13).

**Figure 2.**
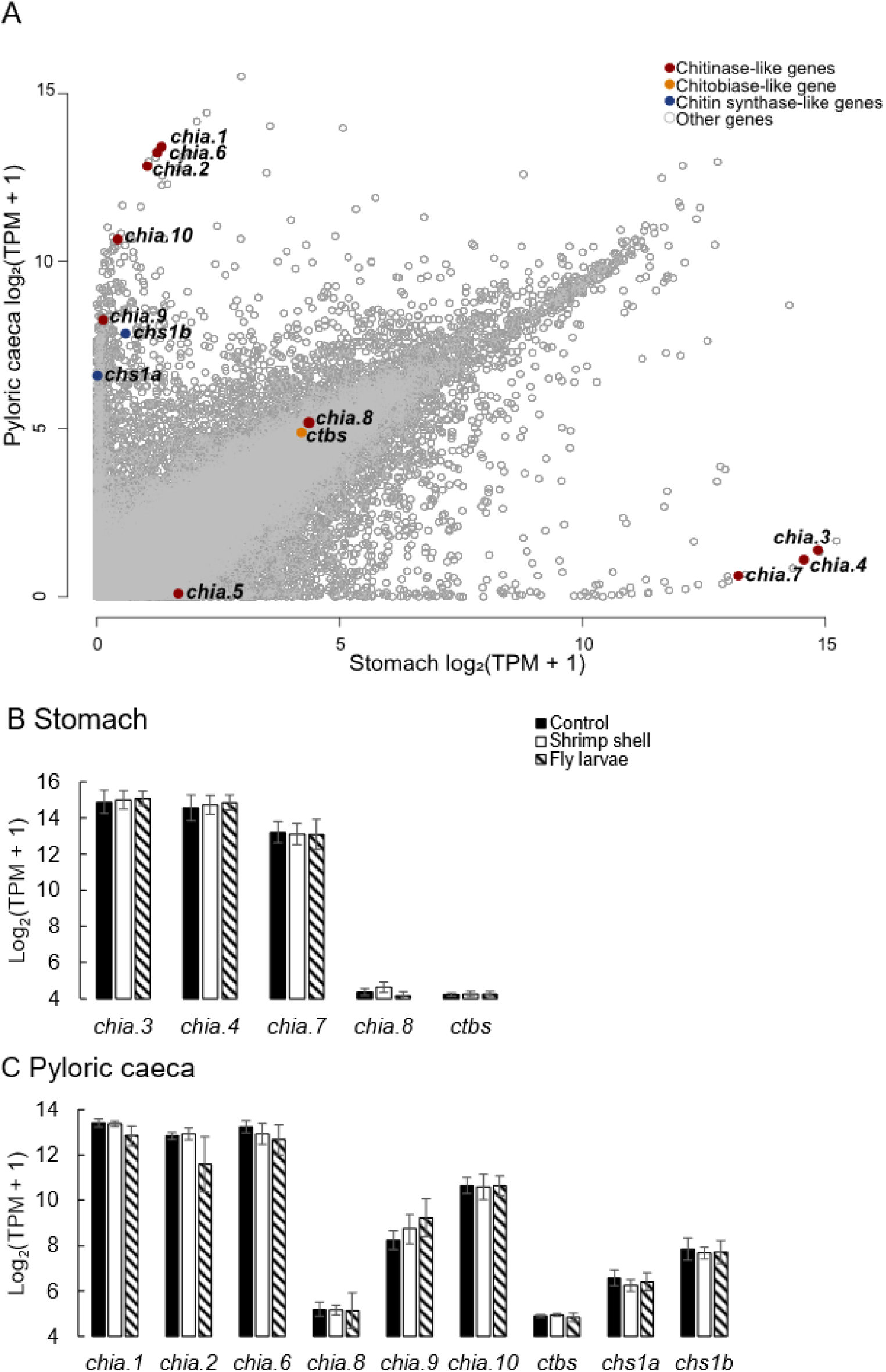
A) Gene expression levels (log_2_(TPM + 1) of chitinase-, chitobiase- and CHS genes compared to the expression levels of all genes in the stomach and pyloric caeca for fish fed the control diet (n=4 for stomach and n=3 for pyloric caeca). B) Gene expression levels (log_2_(TPM + 1) of chitinase and chitobiase genes in the stomach (n=4 for control and fly larvae, n=5 for shrimp shell). C) Gene expression levels (log_2_(TPM + 1) of chitinase and CHS genes in pyloric caeca (n=3 for control and fly larvae, n=5 for shrimp shell). Please note that the y-axis does not extend to 0.

### Gene enrichment analysis

In addition to focusing on genes related to chitin metabolism, we examined how the transcriptome more generally responded to the inclusion of dietary chitin as a replacement to cellulose. Beginning with the stomach, feeding fish a diet including fly larvae had very little effect on the transcriptome compared to the control diet with only one differentially expressed gene (DEG), namely dual-specificity protein phosphatase 1-like (*dusp1*). In contrast, a shrimp shell containing diet provoked a larger effect with 889 upregulated and 570 downregulated genes. The most enriched GO (biological process) and KEGG terms among the upregulated genes were processes and pathways involved in cell organization including muscle structure development, focal adhesion, and extracellular matrix (ECM) receptor interactions (Table 2).

**Table 2.** Top GO terms and KEGG pathways of upregulated (up) and downregulated (down) DEGs in the stomach of fish fed shrimp shells. The rich factor is the ratio of DEG number and the total number of genes annotated to this pathway.

| Enriched term | DEGs | Rich factor | P-value | Method | Mode |
| --- | --- | --- | --- | --- | --- |
| Muscle structure development | 184 | 0.108 | 7.69E-35 | GO_BP | Up |
| Anatomical structure morphogenesis | 396 | 0.063 | 3.18E-27 | GO_BP | Up |
| Cell adhesion | 189 | 0.087 | 5.58E-25 | GO_BP | Up |
| Focal adhesion | 41 | 0.064 | 1.25E-12 | KEGG | Up |
| ECM-receptor interaction | 19 | 0.067 | 8.75E-07 | KEGG | Up |
| Regulation of actin cytoskeleton | 30 | 0.048 | 9.88E-07 | KEGG | Up |
| Synaptic vesicle recycling | 19 | 0.116 | 2.91E-05 | GO_BP | Down |
| Glycoprotein biosynthetic process | 38 | 0.066 | 3.29E-05 | GO_BP | Down |
| Protein glycosylation | 30 | 0.071 | 5.77E-05 | GO_BP | Down |
| UDP-N-acetylglucosamine biosynthetic process | 5 | 0.42 | 3.45E-04 | GO_BP | Down |
| Metabolic pathways | 80 | 0.025 | 4.59E-10 | KEGG | Down |
| Amino sugar and nucleotide sugar metabolism | 11 | 0.085 | 1.20E-06 | KEGG | Down |
| Glutathione metabolism | 9 | 0.084 | 1.20E-05 | KEGG | Down |
| Mucin type O-glycan biosynthesis | 8 | 0.09 | 2.28E-05 | KEGG | Down |

Among the downregulated genes in the stomach, the most enriched processes and pathways were those where glycosylation plays a central role, including synaptic vesicle recycling, glycoprotein biosynthetic processes, and amino sugar and nucleotide sugar metabolism. Notably, several of the downregulated genes are central in the pathway whereby fructose-6-phosphate is converted to uridine diphosphate N-acetylglucosamine (UDP-N-acetylglucosamine biosynthetic process), the substrate molecule used by CHS to synthesize chitin (Figure 3). This includes glutamine-fructose-6-phosphate aminotransferase 2- like (*gfpt2*), glucosamine-phosphate N-acetyltransferase 1 (*gnpnat1*) and UDP-N-acetylhexosamine pyrophosphorylase-like (*uap1*).

**Figure 3.**
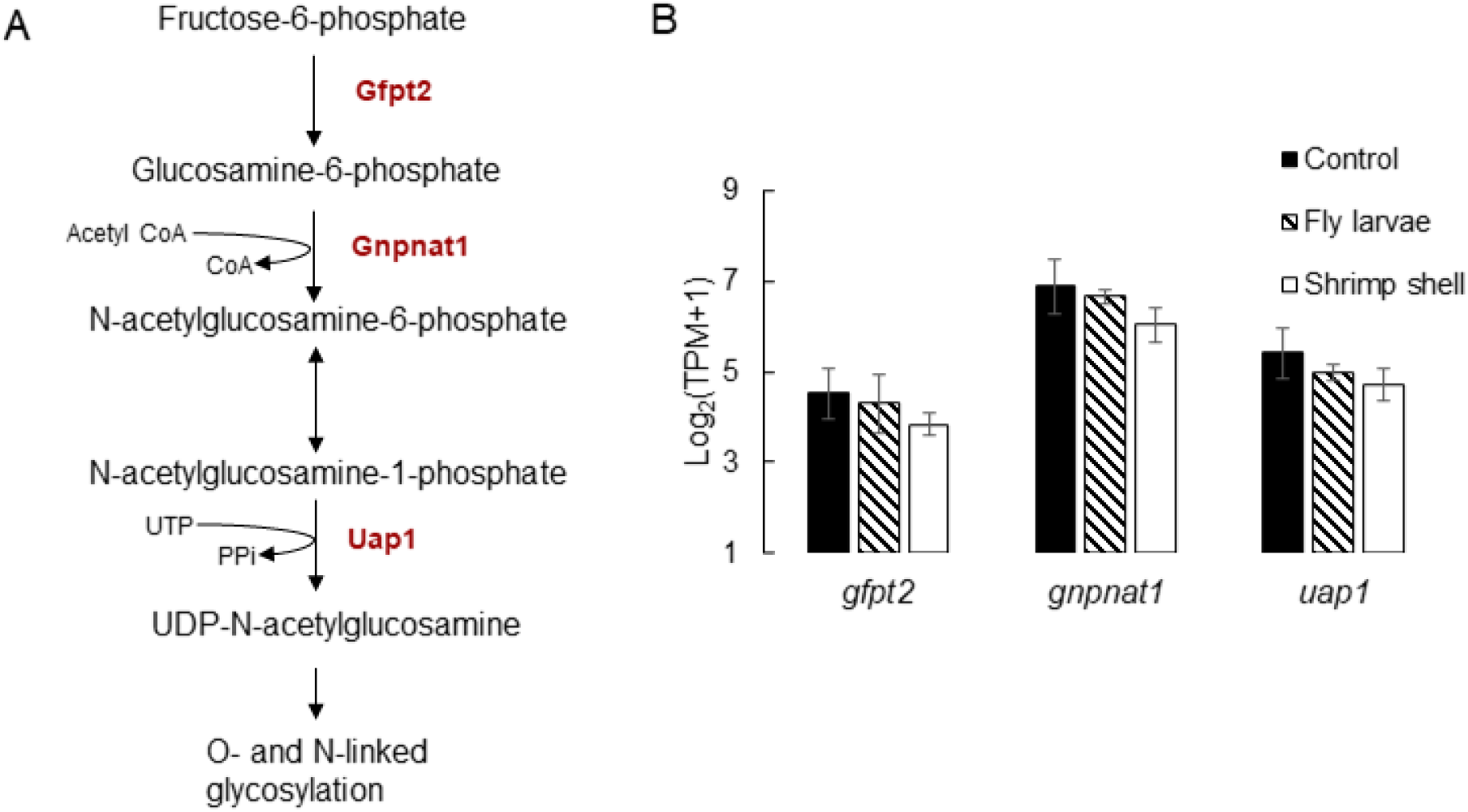
A) The hexosamine biosynthetic pathway leading to the production of UDP-N-acetylglucosamine (UDP-GlcNAc) where central enzymes discussed in the text are marked in red. B) Gene expression levels (log_2_(TPM + 1) of differentially expressed hexosamine biosynthetic pathway genes in the stomach (n=4 for control and fly larvae, n=5 for shrimp shell). Please note that the y-axis does not extend to 0.

When analyzing data from the pyloric caeca only two genes (endonuclease domain-containing 1 protein-like; *endd1*, and ribonuclease P protein subunit p30-like; *rpp30*) were differentially expressed when replacing cellulose with shrimp shell chitin, whereas 53 genes were upregulated and eight downregulated in response to feeding fly larvae. GO enrichment of the 53 upregulated genes revealed that many of these genes (n=22) are involved in the biosynthesis of cholesterol including HMG-CoA reductase (*hmgr*) which is responsible for the rate-limiting step in the cholesterol biosynthesis pathway. No common enriched term was found for the downregulated genes, but the 8 genes detected included acid-sensing (proton gated) ion channel 1 (*asic1*), collagen alpha-1(XXIV) chain-like (*col24a1*), solute carrier family 25 member 48-like (*slc25a48*), peptidase inhibitor R3HDML-like (*r3hdml*), lecithin retinol acyltransferase-like (*lrat*) and three uncharacterized genes.

### The bacterial composition of distal intestine contents

To investigate the impact of dietary chitin on the microbial community, we analyzed the bacterial metapopulation of feed and DI using 16S rRNA gene sequencing. We observed a clear relationship whereby bacteria present in the raw feedstuff were also present in the DI (Figure 4). The microbial profiles of DI contents from fish fed control and shrimp shell diets were almost identical, at the genus level the taxa dominating the distal intestinal contents of fish fed control and shrimp shell were *Lactobacillus* (45.7% and 50.7%), *Streptococcus* (9.4% and 9.3%) and *Weissella* (6.4% and 6.5%). For fish fed fly larvae, the dominant taxa were *Actinomyces* (28.2%), *Bacillus* (21.6%), and *Enterococcus* (16.9%). All prevalent taxa in distal intestinal contents were also present in the feed samples, although the relative amount of *Actinomyces* in fish fed fly larvae was substantially higher than in the feed (28.2% 247 vs 2.18%).

**Figure 4.**
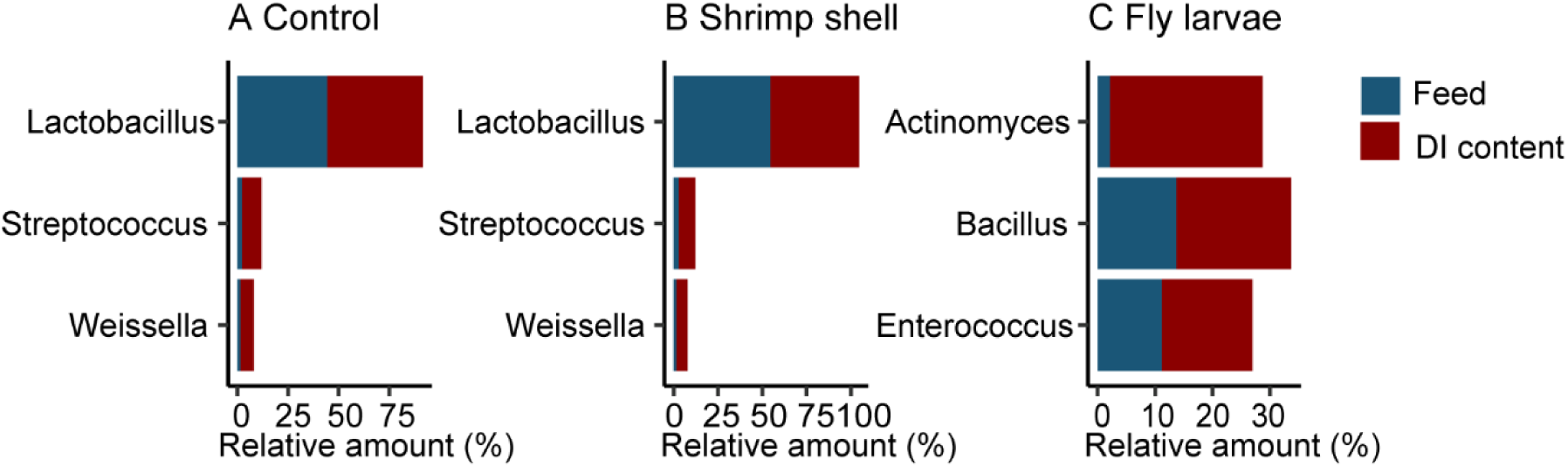
The three most abundant bacteria found in distal intestine (DI content) and fish feed as assessed by 16S rRNA gene sequencing of fish fed A. Control diet (mean value of n=11 for DI content and n=2 for feed), B. Shrimp shell diet (mean value of n=9 for DI content and n=2 for feed), and C. Fly larvae diet (mean value of n=10 for DI content and n=2 for feed).

## DISCUSSION

Transcriptome analyses revealed that the stomach and pyloric caeca react differently to the diets given in this trial. The largest changes in terms of DEGs were observed in the stomach when fish were fed a diet supplemented with chitin from shrimp shells. GO and KEGG analysis lead us to conclude that these changes involve the upregulation of genes involved in tissue organization and extracellular matrix-cell interactions and are linked to structural changes in the gastric tissue. The mucosal barrier that covers the gastrointestinal tract of Atlantic salmon consists of highly glycosylated mucins (Jin *et al*. 2015) and the downregulation of genes involved in glycosylation of these proteins further implies a structural change or a response to the extracellular environment. We hypothesize that this change is partly caused by chitin degradation, with the release of products such as GlcNAc as the result of the activity of highly expressed chitinases in the stomach (Figure 2A-B). In line with what we observe, higher GlcNAc concentrations are associated with the downregulation of genes involved in the biosynthesis of UDP-GlcNAc (Figure 3). UDP-GlcNAc is a substrate required for O – and N-glycosylation of proteins, including mucins that are heavily glycosylated. In mice, orally administered GlcNAc was shown to enter the hexosamine biosynthetic pathway and increase the abundance of UDP-GlcNAc (Ryczko *et al*. 2016). UDP-GlcNAc has shown to act as an end-product inhibitor of this pathway (Chiaradonna *et al*. 2018) and regulates the gene expression of *gfpt2*, which converts fructose-6-phosphate and glutamine to glucosamine-6-phosphate and glutamate, eventually modulating glycosylation homeostasis.

There is evidence for the presence of chitinous structures surrounding the mucosal barrier of ray-finned fish (Tang *et al*. 2015; Nakashima *et al*. 2018) and the results presented here are consistent with the evolutionary conservation of host chitinases and chitobiase to participate in the remodeling of these structures. Previous studies show that Atlantic salmon is not able to utilize chitin to a significant extent (Olsen *et al*. 2006). In line with our results, increasing the dietary chitin content has previously been shown not to correlate with increased chitinase activity in fish (Lindsay *et al*. 1984; Kono *et al*. 1987; Abro *et al*. 2014), and it seems as if the chitinase activity is always present independent of dietary chitin. There could be two possible reasons for this: 1) gut chitinase activity is not regulated by the addition of chitin because Atlantic salmon is exposed to a relatively constant supply of chitin during its life cycle, and/or 2) the chitinase and chitobiase genes are constitutively expressed because of their role as chitin remodelers in the intestinal mucosa. Relatively high CHS gene expression levels in the same intestinal segments of Atlantic salmon as chitinases and chitobiase genes are expressed favor the second hypothesis.

Compared to stomach, few DEGs were detected in pyloric caeca with the greatest changes observed in fish fed fly larvae; this included the significant upregulation of cholesterol biosynthetic genes. Such an upregulation could be expected as a response to lower cholesterol levels in the feed, as insect lipids usually contain low cholesterol levels, but a substantial amount of phytol sterol (Secci *et al*. 2018). This has previously been shown to induce the cholesterol biosynthetic pathway in the pyloric caeca of Atlantic salmon (Jin *et al*. 2018).

Chitinase activity in the stomach and pyloric caeca contents of Atlantic salmon did not seem to be significantly affected by the addition of dietary chitin, but fish fed fly larvae had slightly lower chitinase activity in pyloric caeca contents than fish fed control and shrimp shell diets (Figure 1). This is in line with the slight decrease in gene expression levels of three chitinase genes of fish fed fly larvae; *chia.1, chia.2*, and *chia.6*, all being exclusively expressed in pyloric caeca (Figure 2C). In general, the transcriptome of Atlantic salmon did not seem to respond strongly to changing the standard commercial diet with fly larvae diet, but the bacterial composition of fish fed fly larvae was different from the composition of fish fed control and shrimp shell diets (Figure 4). In accord with our findings, an increase in the relative abundance of *Actinomyces, Bacillus*, and *Enterococcus* when Atlantic salmon is fed fly larvae have previously been reported (Li *et al*. 2021). Since *Actinomyces* and *Bacillus* are potential chitin degraders (Askarian *et al*. 2012; Beier and Bertilsson 2013; Wang *et al*. 2018), we hypothesize that the observed decrease in Atlantic salmon chitinase gene expression levels and activity under fly larvae diet is an effect of the activity of bacterial chitinases, reducing the level of dietary and host-derived chitin available as a substrate to endogenous chitinases in the gastrointestinal tract.

## CONCLUSION

We show that the stomach and pyloric caeca transcriptome of Atlantic salmon did not respond to a great extent to the presence of dietary chitin, in support of the idea that evolutionary conservation of host chitinases is mostly linked to remodeling of chitin as a structural element in the gut lining (Nakashima *et al*. 2018). Furthermore, we demonstrate an association between gut microbial composition, chitin activity in the gut, and host chitinase gene expression, and hypothesize functional interconnection between chitinase-secreting gut bacteria (e.g. *Actinomyces*) and chitinase gene regulation in the host. These results contribute to a greater understanding of chitin metabolism in fish in general.

## DATA AVAILABILITY STATEMENT

Raw RNA-seq data are available on ArrayExpress with the accession number E-MTAB-10594. The raw 16S sequence data are available at NCBI‘s Sequence Read Archive (SRA) with accession number PRJNA820557.

## WELFARE STATEMENT

The experiment was conducted in accordance with Norwegian and European regulations related to animal research. Formal approval of the experiment by the Norwegian Animal Research Authority (NARA) was not required as the experimental conditions were following routine practices at the Centre for Fish Research at NMBU and no compromised welfare was expected.

## ACKNOWLEDGEMENTS

We want to acknowledge Protix Biosystems BV for providing us with insect meal from defatted black soldier fly larvae (*Hermetica illucens*) and Dr. Jon Øvrum Hansen for helping with feed composition calculations and feed pellet processing.

