## Supplementary Table 1 for "The effect of dietary chitin on Atlantic salmon (*Salmo salar*) chitinase activity, gene expression, and microbial composition"

**Supplementary file**

**Supplementary Table 1.** Corresponding NCBI gene IDs to the genes discussed in this paper.

| **Name in paper** | **NCBI gene ID** |
| --- | --- |
| *chia.3* | 106565088 |
| *chia.4* | 106565087 |
| *chia.7* | 106572401 |
| *chia.2* | 106565093 |
| *chia.9* | 106583514 |
| *chia.8* | 106577510 |
| *chia.6* | 106567267 |
| *chia.1* | 106565309 |
| *chia.10* | 106583513 |
| *chia.5* | 106567333 |
| *ctbs* | 100195549 |
| *chs1a* | 106589822 |
| *chs1b* | 106612488 |
| *chs2* | 106562657 |
| *chs3* | 106570137 |
| *gfpt2* | 106611946 |
| *uap1* | 106560290 |
| *gnpnat1* | 100195905 |
